# Genome-scale metabolic modeling of *Ruminiclostridium cellulolyticum*: a microbial cell factory for valorization of lignocellulosic biomass

**DOI:** 10.1101/2025.03.04.641369

**Authors:** Idun Burgos, Ove Øyås, Stéphanie Perret, Henri-Pierre Fierobe, Daniel Machado

## Abstract

The development of sustainable biotechnological processes requires a transition from the traditional fermentation of refined substrates towards the valorization of waste materials such as lignocellulosic biomass. Although these so-called recalcitrant substrates cannot be degraded by model industrial organisms, they can be degraded by microbial consortia through a process of anaerobic digestion, where different community members are able to break down polysaccharides of varied complexity. Among these microbes, *Ruminiclostridium cellulolyticum* stands out as a promising candidate for fermentation of lignocellulose due to its ability to degrade both cellulose and hemicellulose. In this work, we present an updated genome-scale metabolic model for *R. cellulolyticum* strain H10. The model was manually curated with experimental data and the pathways for degradation of cellulose and hemicellulose (arabinoxylan and xyloglucan) were reconstructed and annotated with full detail. The model enables the simulation of the fermentation profile of lignocellulosic materials of various compositions, facilitating the use of this organism as a potential workhorse for sustainable biotechnology, and it provides a valuable template for the reconstruction and optimization of lignocellulose degradation pathways in related organisms.

## Introduction

Anaerobic digestion, in which bacterial communities degrade and ferment biowaste, is a popular approach for biowaste valorization^1^. It is based on a natural process that occurs in, for example, the rumen and in the soil and can yield products like biogas and other valuable fermentation products. However, controlling the anaerobic digestion process is a challenge due to the variability in substrate composition, community structure, inhibitory substances, and metabolic bottlenecks^1^. Due to the intricate nature of the community’s metabolic capabilities and interactions among its members, controlling the carbon flow toward a desired end product, which can range from methane to short or medium-chain carboxylates, remains a challenge^2,3^. This challenge is especially acute given the diversity of metabolic capabilities within the community even though certain functional guilds are conserved^4^. One way to improve control is through microbial community modeling. This approach, based on genome-scale metabolic models (GEMs), has received much attention because of its ability to give mechanistic insights and predict metabolic interactions^5^.

The hydrolysis of cellulose and hemicellulose, the polysaccharide components of lignocellulose, has been shown to be the rate-limiting step or metabolic bottleneck of anaerobic digestion ^6^. There are only a few characterized organisms that have the ability to degrade and use these polysaccharides as a carbon source, due to the recalcitrant structure and complexity of such compounds^7^. One of these organisms is the gram-positive mesophilic bacterium *Ruminiclostridium cellulolyticum*, which has the inherent ability to reduce both hemicellulose and cellulose into their structural components^8^. It produces a multi-enzyme complex called the *cellulosome*, typical of the clostridia growing on crystalline cellulose, which efficiently catalyzes extracellular degradation of lignocellulose. *R. cellulolyticum* is an anaerobic heterotroph with a fermentation profile consisting mainly of acetate, ethanol, lactate, hydrogen, and CO_2_ ^9^. However, as it has a slow metabolism compared to other fermenters, the production rates are typically low. Other cellulolytic clostridia like the thermophile *Clostridium thermocellum* and *Clostridium cellulovorans* have therefore been used as cell factories in similar applications^10^. *R. cellulolyticum* is nonetheless a promising candidate for metabolic engineering due to its almost complete degradation of lignocellulose polysaccharides and a well-characterized cellulosome^11–13^.

The first GEM for *R. cellulolyticum* strain H10, iFS431, was constructed by Salimi et al. (2010)^14^, following the publication of its genome in 2009^15^. The model accounts for 603 metabolites, 621 reactions, and 431 genes. It was built largely based on the annotations available in the JGI database^16^. One of the major limitations of the iFS431 model is that it does not incorporate the pathways for degradation of complex polysaccharides other than cellulose, and the pathway for cellulose degradation is simplified compared to the current knowledge of all the steps involved. This stems from the fact that these pathways have only recently been characterized and annotated in detail^12,13,17–23^. In the meantime, GEMs for other cellulolytic clostridia have also been published. *C. thermocellum* is considered a model species for cellulose-degrading bacteria and has gained much interest for its ability to produce ethanol from cellulose. There are multiple genome-scale metabolic models for this organism^24–26^ with sequential improvements between versions, including more detailed cellulose degradation pathways.

In this work, we present an updated genome-scale metabolic model for *R. cellulolyticum* H10, denoted as iIB728. After an automated genome-based reconstruction, the model was extensively curated based on experimental data obtained from the literature, including growth on multiple substrates, fermentation profiles, cofactor specificity, and mutant phenotypes. In addition, the pathways for degradation and uptake of cellulose, xyloglucan, and arabinoxylan were reconstructed and annotated at an unprecedented level of detail. We demonstrate the applicability of the model in the simulation of batch fermentations using lignocellulosic materials with different compositions. The ability to predict the impact of genetic and environmental perturbations provides a suitable framework to transform *R. cellulolyticum* into a workhorse of industrial biotechnology.

## Results

### Draft model reconstruction

An initial draft reconstruction of *R. cellulolyticum* H10 was created using CarveMe^27^ based on the only available genome of this species in RefSeq (GCF_000022065.1, a high-quality complete assembly). Figure 1 shows a summary of the genes, reactions, and metabolites present in the iIB728 model in comparison with the older iFS431 model. Interestingly, CarveMe automatically reconstructed the pathways for xylan (⍰-1,4 with glucuronic acid decorations), glucomannan, galactomannan, and glucan (⍰-1,3) degradation. These are mostly derived from the *Thermotoga maritima* model iLJ478 ^28^ present in the BiGG database.

**Figure 1.**
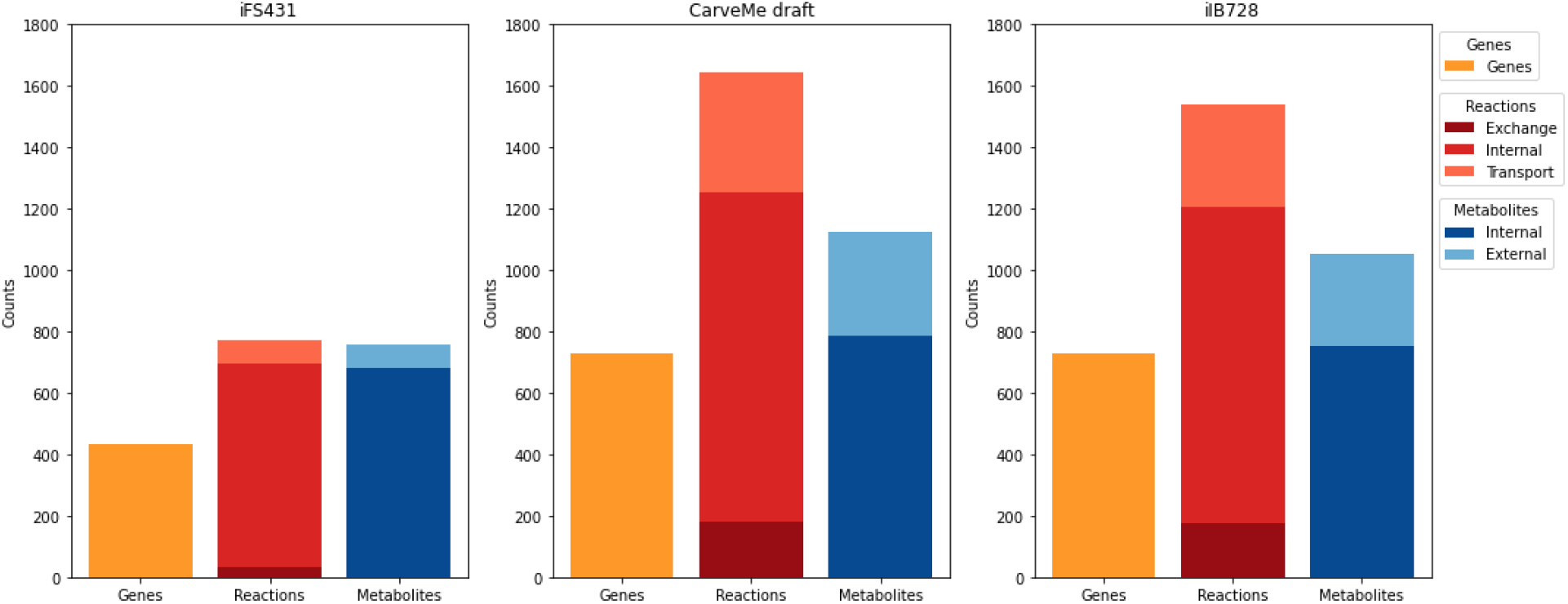
Summary of metabolic models. Total number of genes, reactions, and metabolites are described for the earlier iFS431 model, the initial draft produced with CarveMe, and the final version of the manually curated model, iIB728.

### Manual reconstruction of cellulose and hemicellulose degradation

The most studied pathway for degradation of lignocellulosic components is that of cellulose metabolism. There are extensive studies of both extracellular and intracellular enzymes involved in this pathway^19,21,29,30^. Desvaux (2005) described that oligosaccharides of cellulose, cellodextrins, with a length of up to seven glucose units can be imported and degraded in the cytosol^9^. In recent studies of the ABC transporter for cellodextrins, they reported activity on cellodextrins up to a length of five glucose units^21^ (longer cellodextrins have not been tested, to the best of our knowledge). We reconstructed the transport reactions according to their findings. After being imported into the cytosol by an ABC transporter, the cellodextrins are degraded sequentially by cellobiose/cellodextrin phosphorylases (CEPA). Recent studies have shown that the CEPAs are highly specific to the length of the cellodextrin^19^. The pathway for cellodextrin uptake and degradation was manually reconstructed according to the information above and incorporated into the model (Fig. 2a). All reactions were able to carry flux as long as the corresponding cellodextrin was provided.

**Figure 2:**
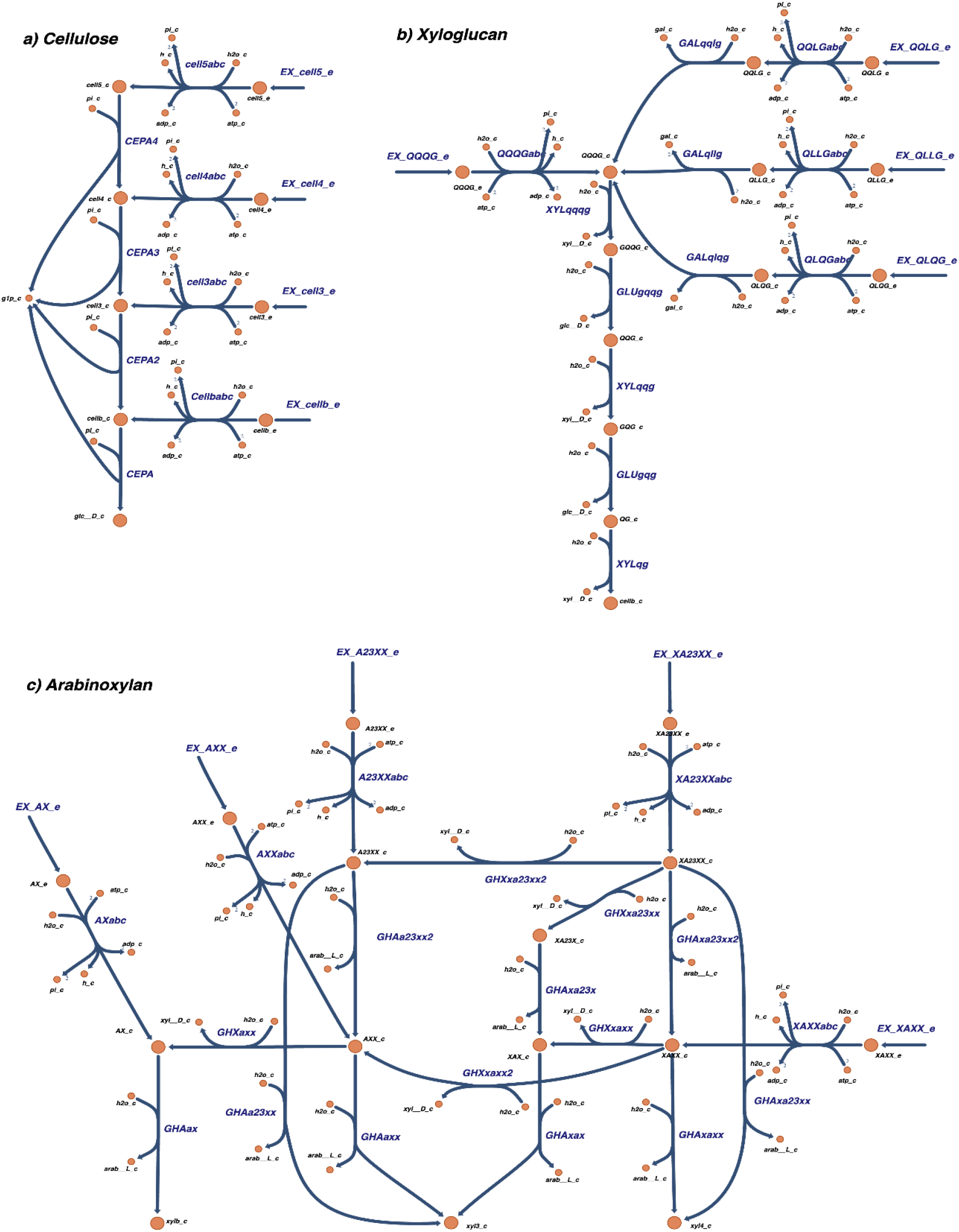
a) Cellulose degradation pathway. This pathway includes exchange reactions for oligosaccharides of different sizes: cellb (cellobiose, two monomer units), cell3, cell4, and cell5 (with 3, 4, and 5 units respectively). Other metabolite abbreviations: glc D (D-glucose), g1p (glucose-1-phosphate), pi (phosphate). **b)** Arabinoxylan degradation pathway. This pathway includes different and highly specific xylanases and arabinofuranosidases. These cut the glycosidic bonds in the main chain composed of ⍰-1,4-linked xylose molecules, and remove the a-1,2- or a-1,3-linked arabinose decorations. **c)** Xyloglucan degradation pathway. This pathway includes a combination of ⍰glucosidases, a-xylosidases, and ⍰-galactosidases that act on different parts of the oligosaccharides. The ⍰ -glucosidases target the main chain of ⍰-1,4-glucose molecules, while the others target the decorations of xylose and galactose.

In addition to cellulose, the metabolism of tamarind xyloglucan has also been extensively studied^17,20,31,31,32^. This metabolism is more complex than that of cellulose due to the composition of this branched heteropolysaccharide. The backbone of xyloglucan is the same as that of cellulose, but the glucosyl units are highly decorated. The xyloglucanases that degrade the polysaccharide extracellularly typically degrade them into oligosaccharides with a glucose backbone length of four units^17^. After transport through an ABC transporter, intracellular degradation of xyloglucan polysaccharides starts with a step-by-step removal of the decorations of galactose and xylose on the glycosyl-residues of the backbone. The first step is the removal of all galactose decorations by beta-galactosidase, followed by the removal of xylose, and then glucose, from the reducing end by alpha-xylosidase and beta-glucosidase, respectively. Using this information, we reconstructed the pathway for xyloglucan degradation and combined it with the model in a similar manner to that of cellulose (Fig. 2b). In the model, the letters G, Q, or L in the metabolite identifiers for oligosaccharides indicate the type of decoration on a glucosyl residue in the backbone. ‘G’ denotes a glucosyl residue without any decoration, ‘Q’ denotes a glucosyl residue decorated with a xylosyl group, and ‘L’ corresponds to a glucosyl residue decorated with a xylosyl group that is further connected to a galactosyl group. In previous literature the ‘Q’ has been described as ‘X’, but we changed this in our model to avoid collision with the identifiers in arabinoxylan (Fig. 1 in Ravachol et al (2016)^17^).

The latest studies of polysaccharide degradation by *R. cellulolyticum* is the degradation of arabinoxylan, deciphered by Liu and colleagues ^23,33^. This included the identification of an ABC importer dedicated to arabinoxylodextrins and its cognate intracellular degradation pathway. Arabinoxylan consists of a D-1,4 xylosyl unit backbone, branched with arabinosyl units (α-1,2 and/or α-1,3). The cellulosomes include xylanases and α-arabinofuranosidases that extracellularly break the polysaccharide into mono- and/or oligosaccharides ^34,35^. However, the extent to which this polysaccharide is really degraded in the extracellular space during the growth of the bacterium remains elusive. Intracellular degradation is catalyzed by two α-L-arabinofuranosidases and two exo-xylanases, which target arabinose decorations and the xylose backbone, respectively^23^. There are also esterases targeting the acetyl, feruloyl and p-coumaroyl decorations^33^, but these were not included in the model as it is not determined how the acetyl groups decorate the xylose backbone. For the α-L-arabinofuranosidases and exo-xylanases we included a reaction in the model for every metabolite following the experimental findings (Fig. 2c). In the model, the letters X and A in the metabolite identifiers for the oligosaccharides indicate the type of decoration on a xylosyl residue in the backbone. ‘X’ represents a xylosyl residue without any decoration, ‘A’ represents a xylosyl with an arabinosyl group. Some xylosyl residues have two arabinosyl groups and the position on the xylosyl residue is given by a number (for example XA2,3XX). In order to avoid gaps, we also included an α-L-arabinofuranosidase active on oligosaccharides XAX and XA2,3X with the respective gene associations. An ABC transporter was also included for every oligosaccharide that the solute binding protein can bind to.

### Manual curation of central carbon metabolism

We aimed to curate the model to perform well under multiple environmental conditions. This is especially important if the model is used in the context of microbial community modeling, where errors in individual models could lead to unrealistic interactions^36^. The first curation step addressed central carbon metabolism. Since *R. cellulolyticum* is a strict anaerobe, the main energy gain occurs through fermentation pathways. Over the years, the species-specific enzymes and cofactors of *R. cellulolyticum* that are involved in glycolysis and fermentation have been studied^29,37,38^. A literature survey was compiled into a dataset of expected phenotypes (see supplementary material). Preliminary simulations using flux balance analysis (FBA) and flux variability analysis (FVA) revealed that the production of the main fermentation products, acetate, ethanol, and lactate, was possible but not growth-coupled. In genome-scale models of fermentative organisms, the way in which cofactors are balanced influences the predicted fermentation profile ^39^. Recent studies have shown that some of the kinases normally driven by ATP are instead driven by GTP or pyrophosphate (PPi) in *R. cellulolyticum* ^20,40^. The same has been shown for *C. thermocellum* ^41^, leading to a predicted increase in ATP yield from glucose in both species ^42^. Furthermore, the fermentation pathways in species of Clostridia are known to be linked to the balancing of NADH, NADPH, and ferredoxin ^43^. We found that some reactions related to ferredoxin balancing were missing in the draft model, including NAD(P)-ferredoxin oxidoreductases, which we manually added.

### Manual curation of carbohydrate transporters

We then used the model to simulate gene deletions and compared the results with experimental data^20^. This resulted in several false positive growth predictions (Table 1), indicating that there were active pathways in the model that were not active or present in the organism, either due to regulatory effects or due to false annotations in the model. By tracing the metabolic flux through the respective pathways, specific reaction targets were removed that would fix the false positive results. We analyzed the evidence for these reactions and their respective proteins by comparing against high-confidence proteins in the UniProt database. This revealed several incorrectly annotated transport reactions in the model, including several phosphotransferase system (PTS) transporters. It has been shown that there are no active PTS transporters in *R. cellulolyticum* ^44^. Removing these reactions resulted in an incomplete pathway for galactose degradation, as the model no longer had a transport system for galactose. Therefore, the PTS transporter for galactose was substituted by an ABC transporter. In addition, UTP-glucose-1-phosphate uridylyltransferase was included to complete the Leloir pathway for galactose metabolism^20^. In addition to these false positive predictions, the model incorrectly predicted that the gene encoding for xylose isomerase (Ccel_3429) was essential for growth in xylose. Kampik et al. (2021) observed that this deletion was not lethal and hypothesized the presence of an alternative enzyme with xylose isomerase activity^20^. Using UniProt we discovered a second gene (Ccel_0500) that is predicted to encode another xylose isomerase. The curated model was able to correctly predict all the experimentally measured mutant phenotypes (Table 1).

**Table 1.** Comparison between experimentally measured phenotypes of single-gene deletions grown on different sugars with model simulations (before and after manual curation) with indication of true positive (TP), true negative (TN), false positive (FP), and false negative (FN) predictions.

| Mutant | Enzyme | Sugar | Growth | Draft model | Curated |
| --- | --- | --- | --- | --- | --- |
| MTL2109 | Cellobiose phosphorylase | Cellobiose | No | FP | TN |
| MTL3221 | Hexokinase | Arabinose | Yes | TP | TP |
| MTL3221 | Hexokinase | Cellobiose | No | FP | TN |
| MTL3221 | Hexokinase | Glucose | No | FP | TN |
| MTL3221 | Hexokinase | Mannose | No | FP | TN |
| MTL3238 | Galactokinase | Arabinose | Yes | TP | TP |
| MTL3238 | Galactokinase | Galactose | No | FP | TN |
| MTL3429 | Xylose isomerase | Arabinose | Yes | TP | TP |
| MTL3429 | Xylose isomerase | Xylose | Yes | FN | TP |
| MTL3431 | Xylulokinase | Arabinose | Yes | TP | TP |
| MTL3431 | Xylulokinase | Xylose | No | TN | TN |

### Calibration of maintenance parameters

The final curation step consisted of calibrating the growth and non-growth-associated maintenance coefficients (GAM and NGAM) using experimental data for chemostat cultivation at multiple dilution rates with cellobiose as the single carbon source^45^ (see Methods). The NGAM value has been previously determined to be 2.2 mmol/gDW/h for growth on cellobiose^14,46^. GAM was found using the same approach as for iFS431, where the sum of squared errors between the experimental and simulated growth rate at increasing uptake rates of cellobiose was minimized (see Methods). We also constrained the production of acetate, lactate, and ethanol to the experimentally determined values for each dilution rate. The GAM value was determined to be 29.54 mmol/gDW (Fig. 3a). This value for GAM is higher than reported for the iFS431 model (15 mmol/gDW) due to differences in the biomass function between the two models. The maximum ATP yield on glucose is higher for iFS431 than iIB728s (Table S1). This difference is accounted for by the presence of a PTS transporter in iFS431, rather than ABC transporter. The use of GTP and PPi instead of ATP in some reactions in the model did not have an effect on the ATP yield, despite what has been suggested previously^20,40^.

**Figure 3.**
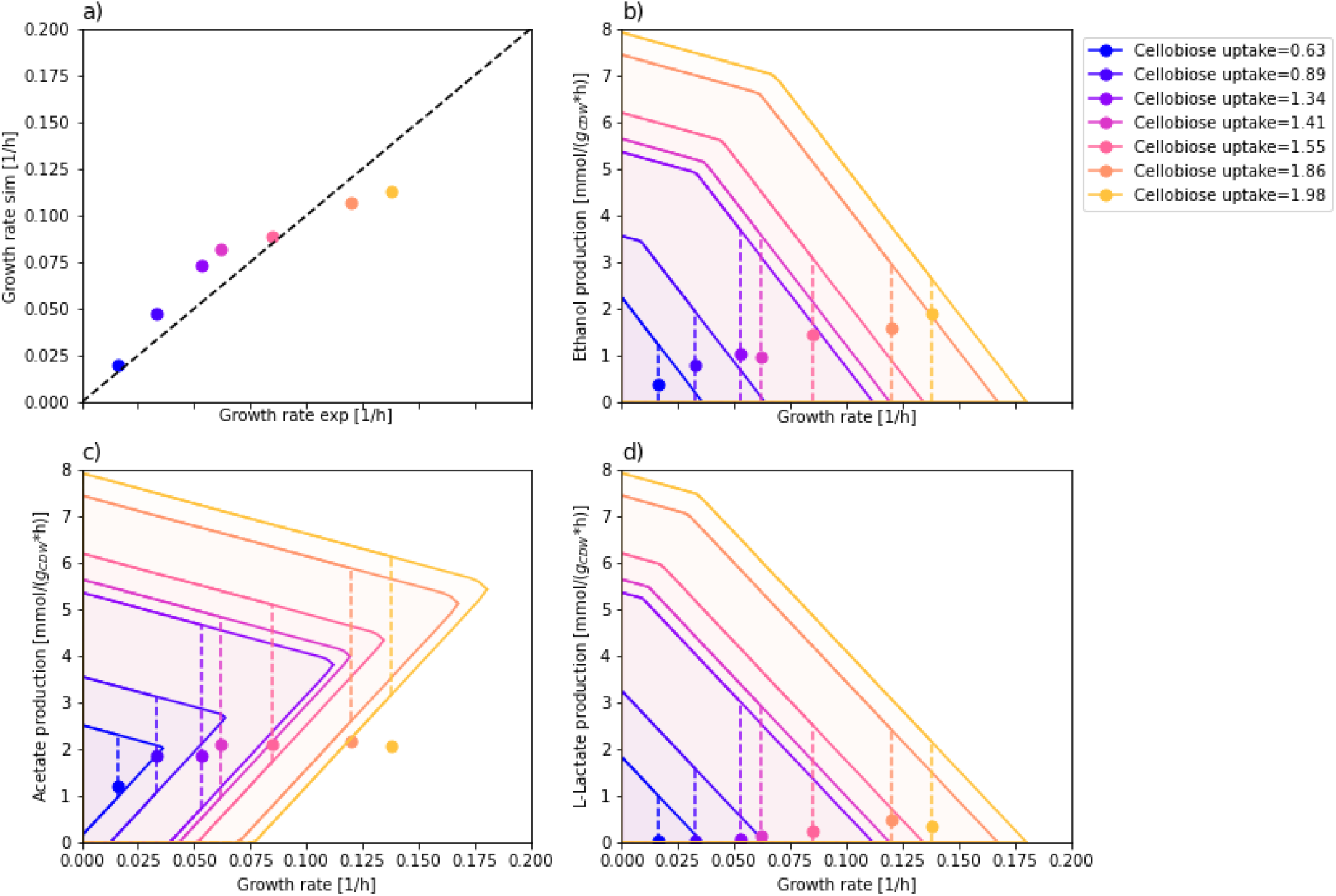
a) Experimental vs simulated growth rates with iIB728 model (when fermentation rates are also constrained. **b-d)** Production envelopes of the main fermentation products. The ranges are obtained with flux variability analysis in a carbon-limited scenario using the experimentally measured cellobiose uptake rate at various dilution rates. The circles indicate the measured production rates of each product at different dilution rates, whereas the dotted lines show the flux variability at each dilution rate.

### Fermentation profile

Using the curated model, we analyzed the profiles of the main fermentation products based on experimental cellobiose uptake rates from Guedon et al. (1999)^45^ (Figs. 3b-d). The updated model predicted growth-coupled acetate production as the main fermentation product, which is in line with previous studies ^45,46^. A strong preference to produce acetate in comparison to ethanol and lactate can be observed, likely because of the increased ATP gain from acetate production. Chemostat experiments show that the organism mainly excretes acetate at lower dilution rates, followed by a steady increase in ethanol production at higher dilution rates, whereas lactate production is quite low, even at higher dilution rates^45^. The experimental production of ethanol and lactate are within the scope of the respective production envelopes and are not growth-coupled in our model. It is believed that there are limitations on specific enzymes around the pyruvate node, like pyruvate-ferredoxin oxidoreductase (PFO)^45^. According to the experimental fermentation profiles, the organism seemingly enters a new metabolic state at one point during the increasing dilution rate. Guedon et al. argued that the metabolic shift occurs due to the oversaturation of pyruvate-ferredoxin oxidoreductase at higher growth rates, causing the flux to be directed toward L-lactate^45^. We can observe that the measured acetate production at the highest growth rate falls outside the computed production envelope (Fig. 4c). This is likely due to an overestimation of the ATP maintenance coefficients, which were fitted to approximate the theoretical and experimental growth rates across all conditions.

**Figure 4.**
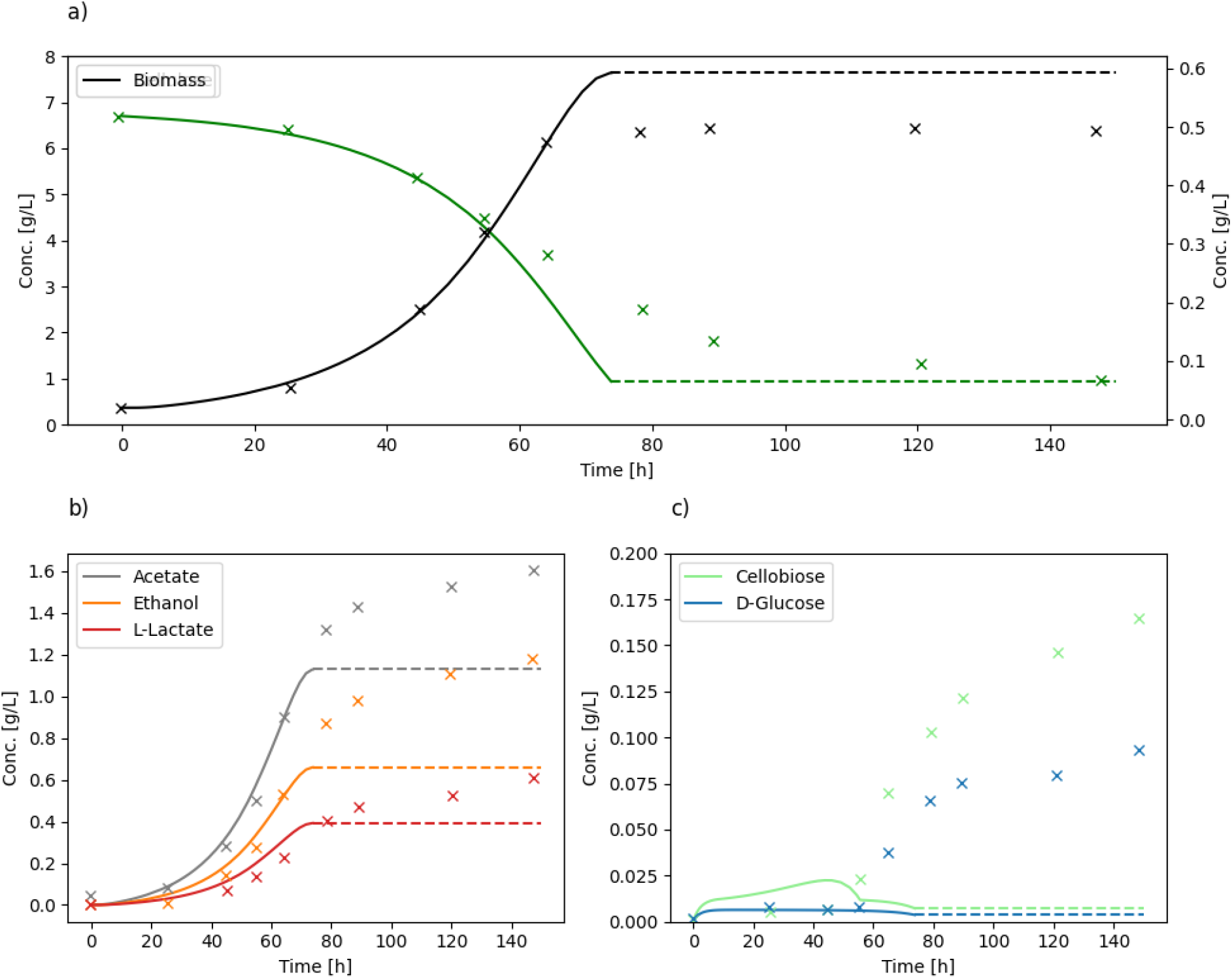
Simulation of batch culture based on experiment by Desvaux et al. (2001) **a)** Cellulose and biomass concentration, **b)** acetate, ethanol and L-lactate concentration, and **c)** cellobiose and D-glucose concentration in the medium. Dotted lines show the final concentrations after the simulation has terminated.

### Simulation of growth on cellulose and hemicellulose substrates

To assess the performance of our model in different conditions we replicated a batch culture simulation on cellulose using dynamic FBA as originally performed by Salimi et al (2010) ^14^. Experimental data for the batch experiment was produced by Desvaux et al. (2001) ^47^. In addition to the starting concentration profiles modeled by Salimi et al (biomass, cellulose, glucose, and cellobiose) we also included flux ratio constraints for the fermentation profiles (see Methods). These were required to force the secretion of the non-growth coupled fermentation products.

The results show that iIB728 is correctly reproducing the uptake, growth, and secretion profiles (Fig. 4). However, similarly to Salimi et al., we were unable to simulate the early termination of growth prior to the depletion of cellulose (Fig. 4a). This is expected because the mechanisms behind the growth termination are not included in the models. This effect is likely caused by the accumulation of toxic metabolites during growth on high concentrations of cellulose, which is a known phenomenon for *R. cellulolyticum* that was also pointed out by Salimi et al. ^9,14^. The experimental results show that product secretion continues, at a lower rate, after growth arrest (Fig. 4b). This is not replicated by the models, since growth (and all metabolic activity) terminates upon substrate depletion. To replicate this effect, we could expand the model with inhibitory terms, but this is outside the scope of this work. We do observe a slightly early arrest of growth before cellulose is fully consumed. This happens due to numerical reasons during simulation when biomass is high and the remaining substrate is not sufficient to fulfill the NGAM requirements for the next integration step.

Overall, the predicted concentration profiles qualitatively fit the experimental results for the period of exponential growth (solid lines in Fig. 4). Additionally, we can observe a low accumulation of glucose and cellobiose, which is also correctly predicted during the exponential phase (Fig. 4c). This likely happens due to the partial binding of the cells to the cellulose, which promotes the immediate use of released sugars and prevents their accumulation in the environment^48^.

One of the main contributions from this work is the incorporation of pathways for cytosolic degradation of oligosaccharides from various mixtures of polysaccharides. To illustrate how this could be used in studies of polysaccharide degradation, we have simulated the growth of *R. cellulolyticum* on wheat straw, which is also a natural source of arabinoxylan. Based on the experimentally determined monosaccharide composition from Patyshakuliyeva et al. (2013)^49^, we estimated the original polysaccharide composition. A previous study showed that *R. cellulolyticum* is able to simultaneously use various simple sugars, although with a preference for glucose^20^. For the degradation of polysaccharides, however, the complexity increases since the structure of the substrate affects the accessibility of the degrading enzymes. A review on ruminal microorganisms uncovered that, in its pure form, hemicellulose is often amorphous and degraded faster than both hemicellulose integrated in the plant cell wall and cellulose ^50^. Therefore, due to lack of suitable data on the kinetics of polysaccharide degradation, we assumed an evenly distributed degradation of polysaccharides with respect to each oligosaccharide (normalized by total carbon content). Figure 5 shows the variation in concentrations of polysaccharides, oligosaccharides, biomass, and fermentation products over an interval of 70 hours. We can observe the simultaneous degradation of the polysaccharides, the accumulation of intermediate oligosaccharides of various sizes, followed by their depletion and secretion of the main fermentation products. This example shows how iIB728 can be used to predict the dynamics of complex polysaccharide degradation, especially as more data becomes available to properly calibrate the kinetic parameters.

**Figure 5.**
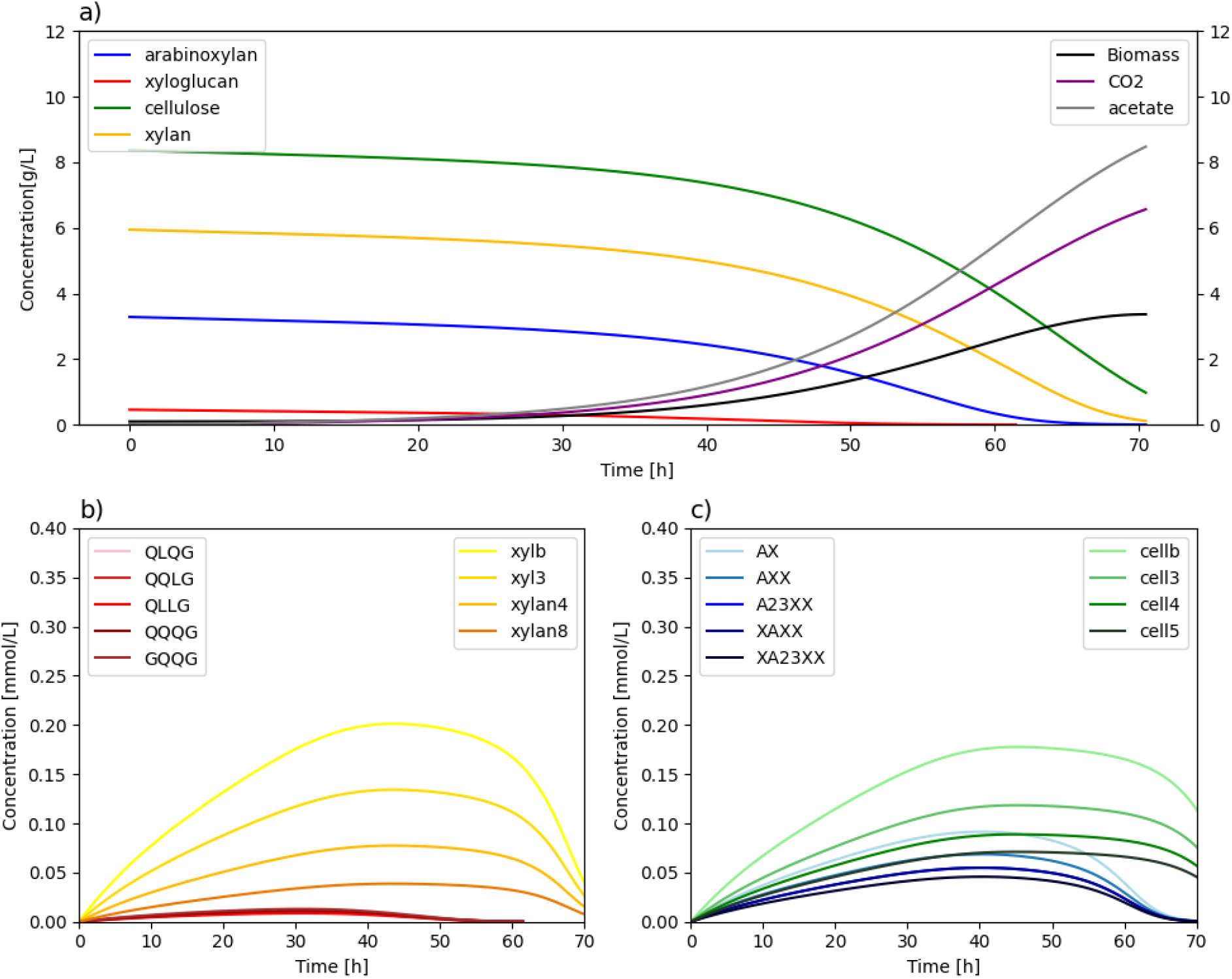
Degradation of a lignocellulosic material. **a)** Concentration of polysaccharides during a simulation of a batch experiment. The left y-axis shows polysaccharide concentration, while the right y-axis shows the biomass, acetate, and CO2 concentration (g/L). **b) and c)** Concentration of multiple intermediate oligosaccharides generated from polysaccharide degradation and gradually consumed. The colors correspond to the original polysaccharide in panel (a), sorted from smaller (lighter) to larger (darker) oligosaccharides.

## Discussion

In this work, we reconstructed a genome-scale metabolic model for *R. cellulolyticum* strain H10 by combining automated reconstruction with manual curation based on experimental data. This model reflects an updated knowledge base aggregating two decades of study of the metabolism of this bacterium. It incorporates a detailed description of the extracellular and intracellular pathways involved in the degradation and transport of lignocellulosic biomass components, including cellulose, xyloglucan, and arabinoxylan.

The initial draft model showed low production of the expected fermentation products. In anaerobic bacteria, cofactor balancing has a major effect on the final product. A common problem in genome-scale metabolic modeling is that cofactor usage cannot be predicted by homology and, in some cases, it is affected by the change of a single amino acid ^51^. There are available methods for predicting cofactor specificity that take into account the protein structure and amino acid residues close to the region of interest ^52–54^. In this case, we took advantage of experimentally determined cofactor dependencies previously reported for ferredoxin, GTP, and PPi^9,20,40^. This alone improved the prediction of fermentation profiles.

It has been previously shown that models generated with CarveMe have lower sensitivity in predicting gene essentiality in comparison to manually curated models, which is not unexpected for a fully automated method. ^27,55^. Despite the small size of the gene deletion dataset used for validation, our results showed a similar pattern with several false positive growth predictions (or, in other words, false negative gene essentiality). This mainly resulted from the automated incorporation of alternative reactions for early steps of sugar catabolism and transport, especially the incorrect inclusion of PTS transporters. Transporter annotation is still one of the main aspects to address for improving automated reconstruction tools^56^. Additionally, the model contained several reactions with more gene associations than reported in the literature. Tracing back the genes to the original model in the BiGG database revealed, in some cases, that the original annotation was also incorrect. The use of manually curated models as basis for annotation can therefore also be a source of inconsistencies for automatic reconstructions.

In this work, we opted to model the degradation of the undigested polysaccharide as an equally distributed composition of oligomers. However, the degradation of polysaccharides depends on where the hydrolyzing enzymes attach to the polysaccharide chain. This is a stochastic process that results in the formation of a pool of oligosaccharides of different sizes. Previous studies have proposed stochastic models for polysaccharide degradation^57,58^, which could be combined with genome-scale metabolic models. Such models could also include the localization of extracellular enzymes, as *R. cellulolyticum* produces both a cellulosome anchored to the cell wall, and free extracellular cellulosomes^59^. This would provide a more realistic description of polysaccharide degradation, although the computational cost might be high and further development of stochastic models is needed, especially for branched polysaccharides. Furthermore, the mixed fermentation profile of *R. cellulolyticum* at higher growth rates indicates that protein constraints are likely at play. It is known that the availability and activity of enzymes is regulated by the concentration of the substrates, cofactors, and downstream products^12,20,22,30,40,47^. Tools like GECKO^60^ can be applied to create a protein-constrained model based on our model. Methods have also recently been developed to create kinetic models from genome-scale metabolic models by network reduction^61^. All in all, there are multiple possibilities to expand the iIB728 model using more detailed modeling frameworks, provided that enough experimental data becomes available for estimation of kinetic parameters.

The main contribution of this study is the reconstruction of pathways involved in the degradation of lignocellulosic oligosaccharides. We have built pathways for the degradation of oligosaccharides from cellulose, xyloglucan, and arabinoxylan. Our model can be used to predict the metabolic phenotype of *R. cellulolyticum* while growing on lignocellulosic substrates with variable composition. Thompson et al. (2016) used the pathway for cellulose degradation in *C. thermocellum* to simulate changes in fermentation profiles on different cellodextrins ^25^. However, this bacterium is not able to grow on hemicellulose ^62^. Therefore, our model provides a valuable template for the reconstruction of other lignocellulose-degrading bacteria obtained from enrichment cultures. Such studies could be pivotal for understanding the role of *R. cellulolyticum* and other microbes in the optimization of the carboxylate platform, paving the way for a stronger bioeconomy based on renewable substrates.

## Methods

### Reconstruction of draft model

The *R. cellulolyticum* strain H10 draft model was produced using CarveMe (version 1.5.1)^27^ with the NCBI RefSeq genome assembly GCF_000022065.1. We used the universal model for gram-positive bacteria and gap-filling using 11 different growth media reported in previous studies ^17,18,20,22,45,46,59^.

### Manual curation steps

The multiple steps for manual curation consisted primarily of: reconstructing the lignocellulose degradation pathways, modifying cofactor specificities according to literature, fixing incorrect gene essentiality predictions, and calibrating GAM/NGAM parameters. For reproducibility, all data extracted from the literature was compiled into spreadsheets and all the model modification steps (adding and/or removing of genes, reactions, and metabolites) are encoded in Jupyter notebooks (available as supplementary material). At each curation step we repeatedly ran a series of tests (using FBA and FVA) to confirm the ability of the organism to grow on the reported growth media and excretes the expected fermentation products, to search for blocked reactions, and to confirm the absence of energy-generating cycles that could be inadvertently introduced. Energy-generating cycles were checked by introducing and maximizing the energy dissipating reactions defined by Fritzmeier et al. (2017)^63^ using parsimonious FBA (pFBA)^64^. This minimizes the number of active reactions, simplifying the process of finding target reactions for curation. For simulations and reconstruction steps both CobraPy (version 0.25.0) and reframed (version 1.2.1) were used.

For the final model, the Python package CobraMod (version 1.3.0)^65^ was applied to add metabolite and reaction annotations based on the MetaNetX cross-reference database^66^. UniProt records were used to add cross-references between gene/protein identifiers in UniProt, RefSeq, and KEGG. Some metabolite charges and chemical compositions were missing, and these were added manually based on entries in the BiGG database^67^. The final version of the model was also tested using the MEMOTE suite^68^.

We calculated GAM through a constrained minimization (bounded between 10 and 50 mmol/gDW/h) of the sum squared error of the differences between experimental and simulated growth rates. The experimental data, taken from chemostat experiments by Guedon et al. (1999)^45^, included growth rates, uptake rate of the limiting substrate (cellobiose), and secretion rates of the main fermentation products. The model was constrained with the rates of the substrate uptake and the fermentation product secretion, and growth rate was maximized.

### Batch growth on cellulose

The modeling framework for the dFBA simulation was based on the dFBAlab implementation in CobraPy^69,70^. We used the same mathematical formulation as given in Salimi et al. (2010)^14^ for cellulose degradation, and glucose and cellobiose uptake rate. In their work, 1 mol of cellulose is broken down to

0.35 mol of cellobiose and 0.3 mol of glucose, which is defined separately from the genome-scale metabolic model. The experimental data used for validation was taken from Desvaux et al. (2001) ^47^ (data from Fig. 2 was digitized using PlotDigitizer^67^). Some changes were made to the method described by Salimi et al. to more accurately reproduce the experimental data. This included lowering the *K*_m_ values for cellobiose and glucose to 0.2 mmol/L, and implementing flux-ratio constraints on the main fermentation products (acetate, ethanol, and L-lactate) based on the experimental measurements. A flux ratio constraint was added between each fermentation product and the biomass reaction. The optimization of growth rate, substrate uptake, and secretion of fermentation products were all handled through the ‘lexicographic constraints’ approach in the dFBAlab implementation, in that specific order.

### Batch growth on different lignocellulose compositions

The modeling framework for the dFBA simulation was based on the dFBAlab implementation in CobraPy, similarly to the previous case study. The kinetic equations used to model our system were as follows:

Specific degradation rate was described using Michaelis-Menten kinetics, with all polysaccharides having the same parameters. The specific degradation rate of a polysaccharide *i, ν*_*poly,i*_, is described by

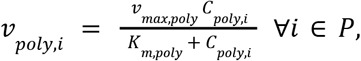

for all polysaccharides in the set of polysaccharides, *P. ν*_*max,poly*_and *K*_*m*.*poly*_ are the parameters for the Michaelis-Menten equation and they are the same for all polysaccharides. *C*_*poly,i*_ is the concentration of polysaccharide *i*. The lower bound for the specific uptake rate of a oligosaccharide *j* from a polysaccharide *i*, 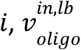, is described by

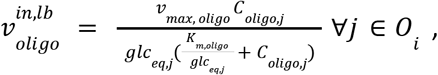

where *O*_*i*_ is the set of oligosaccharides from polysaccharide *i, ν*_*max, oligo*_ and *K*_*m,oligo*_ are the Michaelis-Menten parameters that are the same for all oligosaccharides, and *glc* _*eq,j*_ is the number of glucose equivalents the oligosaccharide amounts to, based on the number of carbon atoms. *C*_*oligo,j*_ is the concentration of the oligosaccharide. As mentioned, 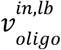 is the lower bound for the oligosaccharide uptake rate. The specific production rate of oligosaccharides, 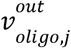, is described by

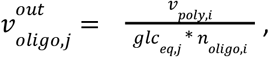

and it is dependent on *ν*_*poly,i*_ as well as *glc*_*eq,j*_ and the number of oligosaccharides for polysaccharide *i, n*_*oligo,i*_ The optimization of growth rate, uptake of oligosaccharides, and secretion of acetate were all handled through the ‘lexicographic constraints’ approach in the dFBAlab implementation, in that specific order.

## Data availability

The curated version of the iIB728 model is publicly available at BioModels.net (MODEL2503030001). All supplementary data and code are publicly available at: https://github.com/IdunBurgos/Ruminiclostridium-cellullolyticum-model-final.

## Acknowledgements

The authors would like to thank the suggestions from Radhakrishnan Mahadevan on the manuscript. This work was supported by project Cell4Chem, funded under the 3rd ERA CoBioTech Joint Transnational Call, with additional support from the Centre of Digital Life Norway (DLN), funded by the Research Council of Norway.

